# Genomic divergence, introduction history and latitudinal adaptation of grass carp

**DOI:** 10.1101/066092

**Authors:** Le Wang, Yubang Shen, Jianjun Fu, Xiaoyan Xu, Gen Hua Yue, Jiale Li

**Affiliations:** Key Laboratory of Exploration and Utilization of Aquatic Genetic Resources, Ministry of Education, Shanghai Ocean University, Shanghai 201306, China; Molecular Population Genetics Group, Temasek Life Sciences Laboratory, 1 Research Link,National University of Singapore, Singapore, 117604, Republic of Singapore; Key Laboratory of Freshwater Fisheries and Germplasm Resources Utilization, Ministry of Agriculture, Freshwater Fisheries Research Center, Chinese Academy of Fishery Sciences, Wuxi 214081; Department of Biological Sciences, National University of Singapore, 14 Science Drive 4, Singapore 117543, Republic of Singapore; School of Biological Sciences, Nanyang Technological University, 60 Nanyang Drive, Singapore 637551, Republic of Singapore

**Author notes:** Corresponding author; Dr. Gen Hua Yue Temasek Life Sciences Laboratory, 1 Research Link, National University of Singapore, 117604 Singapore. Dr. Jiale Li Key Laboratory of Exploration and Utilization of Aquatic Genetic Resources, Ministry of Education, Shanghai Ocean University, Shanghai 201306, China. Le Wang and Yubang Shen contributed equally to this study. NGS data was deposited in the DDBJ Sequence Read Archive (Project accession no. PRJDB4785).

**Keywords:** grass carp, genomic divergence, introduction, local selection, latitudinal variation

## Abstract

Understanding the genomic signatures of population differentiation is fundamental to obtain a comprehensive view of the evolutionary process of organisms. Grass carp is one of the most important fish species in the world due to its significant value in aquaculture and world-wide vegetation biocontrol. However, little is known about the contemporary population structure and also the genetic basis of adaptation to a wide range of latitudinal environments. Using 43310 SNPs generated by genotyping by sequencing in 197 grass carps from nine populations, we examined the patterns of population differentiation, historical introduction and evidence of local selection. The overall genetic differentiation across all native populations was unexpectedly low. Nevertheless, these native populations were clearly differentiated into three genetic clusters, corresponding to the Yangtze River, the Pearl River and the Heilongjiang River System, respectively. Populations in Malaysia, India and Nepal, with the earliest introduction records, most likely have an origin from the Pearl River System. Using conceptually different approaches, 451 loci were detected under potential local selection, among which 84 were annotated to have a gene feature. 19.0% of the genes under putative selection were involved in immune responses, while 42.9% of the annotated loci showed a signature of latitudinal variation. This study provides valuable information for application of genomic tools in addressing questions concerning population differentiation that was influenced by both neutral and adaptive forces, as well as human activities.

## Introduction

Grass carp (*Ctenopharyngodon idella*), belonging to the family Cyprinidae, is a large herbivorous freshwater fish species (Froese and Pauly 2015). It is of great importance as a food fish species as well as a species for world-wide aquatic vegetation control (Cross 1969; Lembi *et al.* 1978). Grass carp is native to eastern Asia and broadly distributed from the Heilongjiang River System (Amur River) southward to northern Vietnam (Froese and Pauly 2015). According to literature records, grass carp has a culture history of more than 1300 years since the Tang Dynasty (FAO 2014). The aquaculture practices are mainly conducted within the geographical regions of the Yangtze and the Pearl River Systems of China (FAO 2014). Recent success in artificial breeding has significantly promoted the aquaculture industry of this species (Stanley 1976; Boney *et al.* 1984; Allen Jr and Wattendorf 1987; Peter *et al.* 1988). The annual global production has been over 5 million tons since 2013 with an estimated economic value of 5 billion US dollar (FAO 2014). Grass carp is of the highest production yield among all the farmed fish species around the world and accounts for approximately 15.6% of global freshwater aquaculture production (FAO 2014).

Due to herbivorous habits, grass carp has been broadly introduced to more than 40 countries around the world to control the undesirable and/or invasive aquatic plants of freshwater systems (Skelton 2001). Artificial introductions were intensively conducted since the 1960s (Welcomme 1988). The earliest introduction of grass carp was documented from Southern China to Malaysia by Chinese immigrants in the 1800s (Welcomme 1988). However, which native populations these introduced grass carp originated from is unclear. Recently, many studies have reported that introduced/invasive grass carp have endangered native ecological systems and caused great economic loss because they can completely eliminate vegetation from freshwater systems, destroy the populations of native fish species and introduce parasites (Moyle 1986; Chilton II and Muoneke 1992; Bain 1993).

Within its native distribution range, grass carp mainly lives in three independent river systems: the Heilongjiang River, the Yangtze River and the Pearl River System (Fu *et al.* 2013). Understanding range-wide population structure is critical to conserve and utilize the genetic resources (Avise 1992). However, it is still not clear, due to a lack of genetic studies (Zhang *et al.* 2001; Liu *et al.* 2009; Fu *et al.* 2013). In particular, grass carp has been cultured for more than 1300 years (FAO 2014). It is also not known if aquaculture practices with such long history have left significant imprints on the contemporary population structure. Importantly, a geographical pattern of population differentiation is the genetic basis to trace the introduction from native habitats to foreign environments (Cornuet *et al.* 1999; Paetkau *et al.* 2004). Investigation on the environments of both native and foreign habitats can provide critical biological information for setting up effective introduction plans (Lande 1988; Bain 1993). It can also mitigate the adverse effects on the foreign habitats that were caused by grass carp as an agent of biological invasion (Cross 1969; Bain 1993).

Local selection is of critical importance in the evolution of species (Savolainen *et al.* 2013). Genetic studies focusing on different environments can provide crucial information for understanding the selective forces driving local adaptation (Sultan and Spencer 2002). In some cases, however, local selection acts along certain geographical gradients, e.g. latitude, longitude and altitude, and shows the same direction as the overall neutral forces (Storz 2002; Vasemägi 2006). Thus, it is more challenging to discriminate adaptive evolutionary forces from background neutral forces (McKay and Latta 2002; Storz 2002). Nevertheless, these environmental factors are of great interest in studying the selective forces that shape adaptive divergence (Gilchrist and Partridge 1999; Alberto *et al.* 2013). Significant associations between environmental variables and genetic markers are typically investigated and considered as footprints of local selection (Sezgin *et al.* 2004; Vasemägi 2006; Antoniazza *et al.* 2010). Although adaptive variation can be consistent with “isolation-by-distance” in geographical pattern and also explained by neutral forces like gene flow, genetic drift and admixture between adjacent populations, the footprints of local selection still can be inferred by comparing the relative pattern and strength of population differentiation against environmental variables between candidate loci of adaptive divergence and neutral markers (Storz 2002; McKown *et al.* 2014).

Grass carp are naturally distributed in a wide range from Southern Siberia to northern Vietnam spanning approximately 30 latitudinal degrees and is tolerant of extreme temperature from 0° to 38°C (Froese and Pauly 2015). The distribution is much likely limited by habitat temperature, and even correlated to the latitudinal variation of temperature. Due to the small number of markers, e.g. microsatellites, however, previous studies only focused on identifying the neutral genetic variations and population structure (Zhang *et al.* 2001; Liu *et al.* 2009; Fu *et al.* 2013). The lack of useful high-density genetic markers limited our understanding on adaptive population differentiation, particularly along latitude (Narum *et al.* 2013).

Here, a total of 197 grass carp, including both wild populations throughout the whole distribution range and populations with the earliest introduction history were analyzed using ddRAD-Seq approach (Peterson *et al.* 2012). First, the aim was to identify range-wide population structure and examine the pattern of gene flow, which can help understand the geographic and demographic factors that have played crucial roles in population differentiation of grass carp. Furthermore, we intended to test if the level of population differentiation is sufficient to trace the introduction of grass carp by using the populations with the earliest introduction history. The results can provide useful information for tracing global introduction of grass carp and better utilizing this species in the biocontrol of aquatic vegetation. In addition, as the native populations show a significant latitudinal distribution pattern, our goal was to detect the footprints of local selection and discuss if such divergence was correlated to specific environmental factors, particularly temperature. Finally, we aimed to identify candidate genes under potential directional selection, which help understand the mechanism of evolution under selection. In total, this study can disentangle the effects of demographic history, gene flow and local selection on the contemporary population differentiation and provide important information for both utilizing this species as a tool in biocontrol and understanding the adaptive divergence of freshwater fish species in the presence of complicated gene flow and demographic history.

## Materials and Methods

### Sampling and data collection

Grass carp including six wild and three introduced populations consisting of 197 individuals were collected between 2007 and 2008. The wild populations were from the three river systems: the Heilongjiang River, the Yangtze River and the Pearl River System, across this species’ distribution range, while the three introduced populations were sampled from Malaysia, India and Nepal, respectively (Table 1 & Figure 1). The annual average temperature of each sampling site was retrieved from weather.sina.com.cn (Table 1). Population Malaysia was documented as being introduced in the1800s from southern China, while population India was recorded as being introduced from Hong Kong, China (the Pearl River System), in 1959 and 1968 (Welcomme 1988). Population Nepal was set up by introduction from India in 1966-1967 (Shireman and Smith 1983). All samples were estimated as more than one year old. Fin tissue was collected and preserved in 95% ethanol at -20°C. Genomic DNA was isolated using DNeasy Blood & Tissue Kit (Qiagen, Germany) and quantified using Qubit^®^ assays (Life Technologies, USA).

**Table 1.** Sampling information of six native and three introduced grass carp populations including river systems of origin, numbers of samples, sampling localities and dates, and the annual average temperature of each sampling locality. Measures of genetic diversity including observed heterozygosity (*H_O_*), expected heterozygosity (*H_E_*) and nucleotide diversity (π) are also indicated.

| Samples | Origin | N | Longitude | Latitude | Date | Temperature | $H_O$ | $H_E$ | $\Pi$ |
| --- | --- | --- | --- | --- | --- | --- | --- | --- | --- |
| Nenjiang | Heilongjiang River System | 22 | 125.22 | 49.21 | 2007 | 4.3 | 0.197 | 0.202 | 0.207 |
| Hanjiang | Yangtze River System | 26 | 119.43 | 32.35 | 2007 | 15.4 | 0.200 | 0.201 | 0.205 |
| Jiujiang | Yangtze River System | 23 | 115.96 | 29.72 | 2007 | 17.7 | 0.202 | 0.205 | 0.210 |
| Shishou | Yangtze River System | 11 | 112.39 | 29.74 | 2007 | 16.9 | 0.210 | 0.204 | 0.214 |
| Zhaoqing | Pearl River System | 21 | 112.53 | 23.08 | 2007 | 22.7 | 0.203 | 0.201 | 0.206 |
| Vietnam | Pearl River System | 26 | 105.98 | 21.12 | 2008 | 24.8 | 0.190 | 0.191 | 0.195 |
| Malaysia | Introduced | 18 | 101.15 | 4.58 | 2008 | Na. | 0.136 | 0.126 | 0.130 |
| India | Introduced | 25 | 83.37 | 26.76 | 2008 | Na. | 0.171 | 0.168 | 0.171 |
| Nepal | Introduced | 25 | 85.03 | 27.42 | 2008 | Na. | 0.161 | 0.152 | 0.155 |

**Figure 1.**
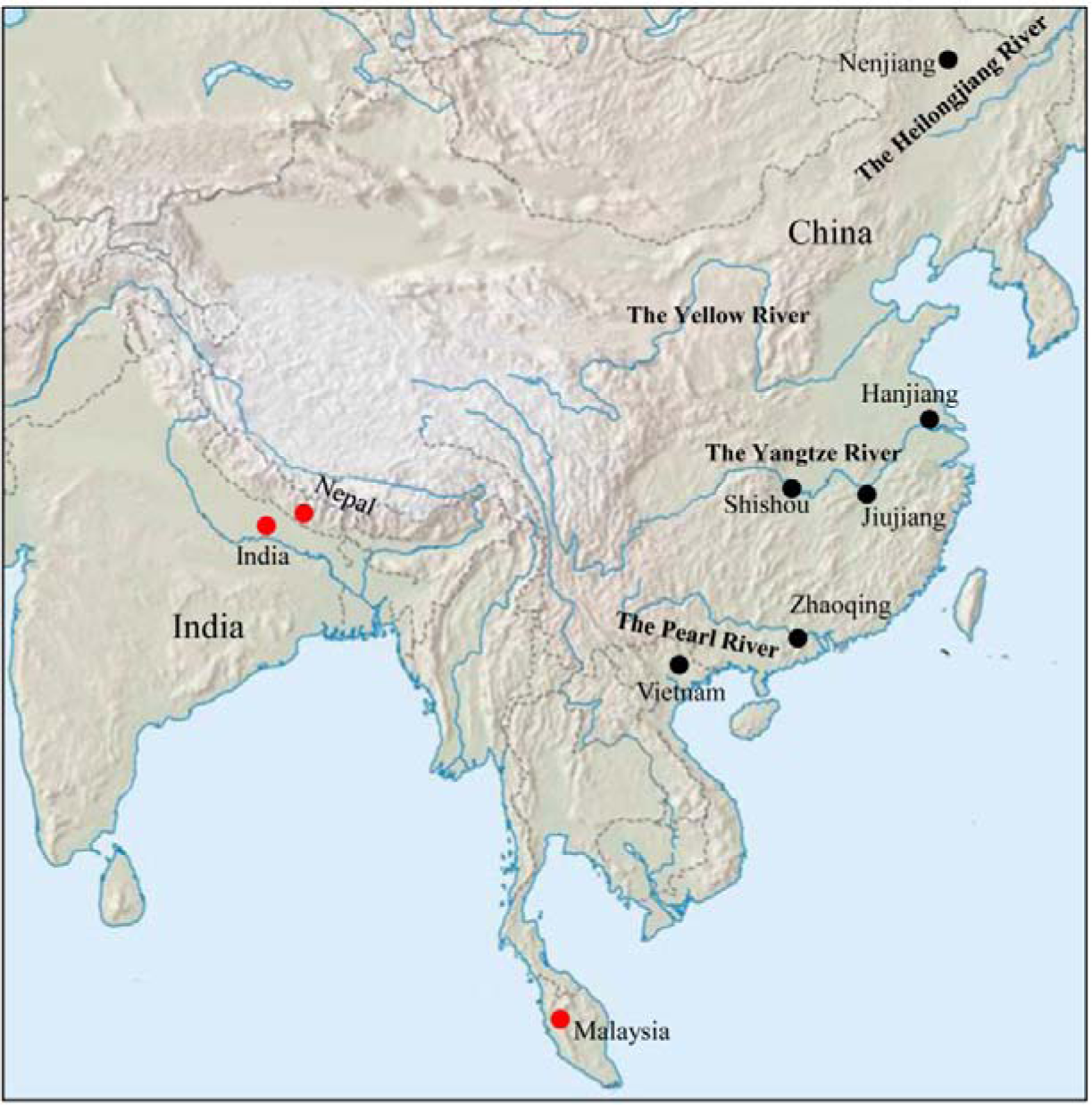
Figure 1 Sampling sites of six native grass carp populations distributed in the three river systems: the Heilongjiang River, the Yangtze River and the Pearl River Systems, and three introduced populations from Malaysia, India and Nepal. The wild and introduced populations are denoted as black and red solid circles, respectively. Detailed sampling information is listed in Table 1.

### enotyping by sequencing

Genotyping by sequencing (GBS) was conducted using the ddRAD-Seq approach (Peterson *et al.* 2012). Restriction enzymes PstI-HF and MspI (New England Biolabs, USA) were selected for library construction. 200 ng genomic DNA was fully digested with two enzymes. Digested fragments were ligated with barcoded adaptors using T4 ligase (New England Biolabs, USA) and then pooled for cleanup with QIAquick PCR Purification Kit (Qiagen, Germany). The cleaned products were size selected and purified (300-500 bp) by running gels and using QIAquick Gel Extraction Kit (Qiagen, Germany), respectively. The recovered libraries were then amplified using Phusion^®^ High-Fidelity DNA Polymerase (New England Biolabs, USA). After a final cleanup using QIAquick PCR Purification Kit (Qiagen, Germany), the libraries were sent to a NextSeq 500 platform (Illumina, USA) for 2x150 bp paired-end sequencing.

The program *process_radtags* (Catchen *et al.* 2011) was employed to filter the raw sequencing reads with default parameters and reads with any uncalled base were removed. Clean reads were then demultiplexed and trimmed to 100 bp for *in silico* mapping. First, reads were mapped to the reference genome of grass carp v1.0 (Wang *et al.* 2015b) using the program BWA-MEM with default parameters (Li and Durbin 2010). Reads with multiple targets in the reference were excluded from further analysis. Reference-aligned reads were then assembled into stacks for each individual using *pstacks* implemented in the package Stacks v1.34 (Catchen *et al.* 2011). A total of 54 individuals randomly selected from each population were used to construct a catalogue of stacks using *cstacks*. Stacks from each individual were then matched against the catalogue for SNP discovery using *sstacks*. Finally, genotyping was conducted across all populations using the program *populations* with a minimum of 10× sequence depth. SNPs were further filtered to meet the following criteria: present in > 70% of the individuals in each population, have no more than two alleles and show an observed heterozygosity of < 0.5 (Hohenlohe *et al.* 2010). Only one SNP was retained for each RAD locus. Hardy-Weinberg equilibrium (HWE) for each locus was examined using Genepop v4.2 (Raymond and Rousset 1995) and loci that deviated from HWE in a single population at the significance level of 0.01 were excluded from further analysis.

### Population structure and phylogenetic relationship

Genetic diversity for each population was measured by observed heterozygosity (*H_O_*), expected heterozygosity (*H_E_*) and nucleotide diversity (π), while genetic divergence for each individual locus was estimated using *F-statistics* (Weir and Cockerham 1984). All these calculations were performed using the program *populations* (Catchen *et al.* 2011). Population genetic divergence was estimated in the form of pairwise *F_ST_* using the program Arlequin 3.5 (Excoffier and Lischer 2010). Statistical significance was examined using an exact test with 10 000 permutations. Population structure at both the population and individual levels was investigated by principle component analysis using the program Eigenstrat v5.1 (Price *et al.* 2006). The pattern of population differentiation was examined in the form of isolation-by-distance (IBD) using Mantel tests with the program IBD v1.52 (Bohonak 2002). The genetic distance was measured using *F_ST_*/1-*F_ST_*, while the geographical distance was estimated as the linear distance between sampling localities.

The phylogenetic relationship among populations was constructed using a Neighbor-Joining approach with the program Populations v1.2.33 (Langella 1999) by bootstrapping over loci for 1000 times. The origins of introduced populations from native populations were determined using ancestral alleles. The software fastStructure v1.0 (Raj *et al.* 2014) was employed to infer the ancestral alleles between one introduced population and two native populations. The two selected native populations were from the Yangtze River and the Pearl River systems, respectively, and had the closest phylogenetic relationship with the introduced population. The program was run 10 times for each K value (from 1 to 6) with default parameters. The most likely number of genetic clusters (K) was estimated by plotting the marginal likelihood value.

### Identifying footprints of selection

In order to identify evidence of latitudinal variation at SNP loci, we independently estimated the association between allele frequencies for each SNP and latitude at the population level using a liner correlation method. Pairwise genetic distance based on allele frequencies of individual locus was estimated according to the method by Reynolds *et al.* (1983). Genetic distance was then correlated to geographical distance among populations using Mantel tests to discriminate neutral mutations from the loci showing latitudinal variation in allele frequencies. By removing the loci under putative neutral processes, a set of candidate SNPs showing latitudinal variation but not isolation-by-distance in genetic divergence was obtained.

Evidence of local adaptation was detected for individual locus using a Bayesian generalized linear mixed model involving covariance of allele frequencies and environmental variables with the program Bayenv (Coop *et al.* 2010). A Bayes factor (BF) was calculated for each SNP to measure the strength of the correlation between SNP variation and environmental variables. According to the method by Coop *et al.* (2010), a BF > 3 was considered as a substantial evidence for selection. The program was run for five times with an independent variance-covariance matrix of population genetic variation to achieve consistency among the runs.

*F_ST_*-based outlier tests were also performed to identify signatures of spatial purifying selection. Outlier loci under directional selection are expected to show higher levels of divergence, while loci under balancing selection would show lower levels of genetic divergence compared to the putative neutral loci (Beaumont and Nichols 1996; Foll and Gaggiotti 2008). Firstly, a Bayesian simulation-based test implemented in BayeScan (Foll and Gaggiotti 2008) was used to identify outlier SNPs. Loci with Bayes factor > 3 were considered as substantial outliers. Considering that grass carp are distributed in different river systems and that there is among-group genetic structure, a hierarchical island model (Beaumont and Nichols 1996) for identifying outlier loci was also employed using the program Arlequin 3.5 (Excoffier and Lischer 2010) with the following parameters: 50 000 simulations, 10 simulated groups, and 100 demes per group. Only outliers above the 99% quantile of the null distribution were considered as candidates under spatial purifying selection.

### Analysis of the genes under putative selection

Loci under putative directional selection were functionally annotated by Blast2Go (Conesa *et al.* 2005) against all available nucleotide databases with an E-value cutoff of 10-6. SNPs within both exons and introns were considered to have a gene feature. Enrichment of Gene Ontology (GO) terms was conducted using the program WEGO (Ye *et al.* 2006) with default parameters. Loci were also mapped to the reference genome of zebrafish (GRCz10) using Blastx to retrieve the corresponding Ensembl gene IDs. A more detailed functional annotation of these genes were then performed by mapping to the Kyoto Encyclopedia of Genes and Genomes (KEGG) pathway database (Kanehisa and Goto 2000) using the program David (Huang *et al.* 2009). The signaling pathways of at least two genes in default were enriched for further analysis.

## Results

### SNP discovery and genotyping

In total, NGS produced an average of 10.17 million raw reads for each individual. After quality control, 9.21 million reads per individual were obtained for sequence mapping and SNP discovery. A total of 280544 SNPs were discovered across all nine populations. After removing the loci that failed to meet the filtering criteria, 43310 SNPs were genotyped across all populations, among which 35844 (82.8%) showed minor allele frequency (MAF) of > 0.05.

### Genetic diversity and population structure

The measures of genetic diversity, including *H_O_*, *H_E_* and π, estimated based on all variant SNPs were shown in Table 1. For the six wild populations, Vietnam showed slightly lower genetic diversity than the others. We also observed that the genetic diversity of the introduced populations including Malaysia, India and Nepal were significantly lower than that of the wild populations (*P* < 0.001, one-way ANOVA test).

Pairwise *F_ST_* analysis revealed significant genetic differentiation between the wild populations and the introduced populations with *F_ST_* ranging from 0.1126 between Zhaoqing and India to 0.2399 between Vietnam and Malaysia (Table 2). However, genetic differentiation between each pair of wild populations was shallow with *F_ST_* ranging from 0.0073 between Jiujiang and Shishou to 0.0515 between Hanjiang and Vietnam, although significantly different from 0. The wild population Zhaoqing from the Pearl River System showed slightly lower genetic differentiation with the three introduced populations: Malaysia, India and Nepal, compared to the other wild populations (P < 0.05, paired *t-test*). For wild populations, genetic divergence at individual locus was estimated between the most divergent populations (Hanjiang vs Vietnam) and also between the most distant populations (Nenjiang vs Vietnam). We observed that no loci showed *F_ST_* > 0.5 and only < 15 % of loci had *F_ST_* > 0.1 for both population pairs (Figure 2). Loci showing *F_ST_* > 0.1 were found more frequently between Nenjiang and Vietnam (14.5%) than between Hanjiang and Vietnam (5.6%). Principle component analysis revealed that the wild populations were strikingly differentiated from the introduced populations (Figure 3a). For the wild populations, Zhaoqing and Vietnam from the Pearl River System were clearly differentiated from the populations from both the Yangtze River System (Hanjiang, Jiujiang and Shishou) and the Heilongjiang River System (Nenjiang), although there was some mixture of individuals between the Yangtze River and the Heilongjiang River Systems (Figure 3b). The pattern of genetic differentiation across all wild populations rejected the model of isolation-by-distance (*R*^2^ = 0.187, *P* = 0.16; Figure 4a). Considering that some individuals of the Heilongjiang River System very likely have an origin from the Yangtze River System (Figure 3b), the population Nenjiang was further removed from Mantel tests. Interestingly, we identified a strong correlation between population differentiation and geographical distance (*R*^2^ = 0.876, *P* < 0.001; Figure 4b).

**Figure 2.**
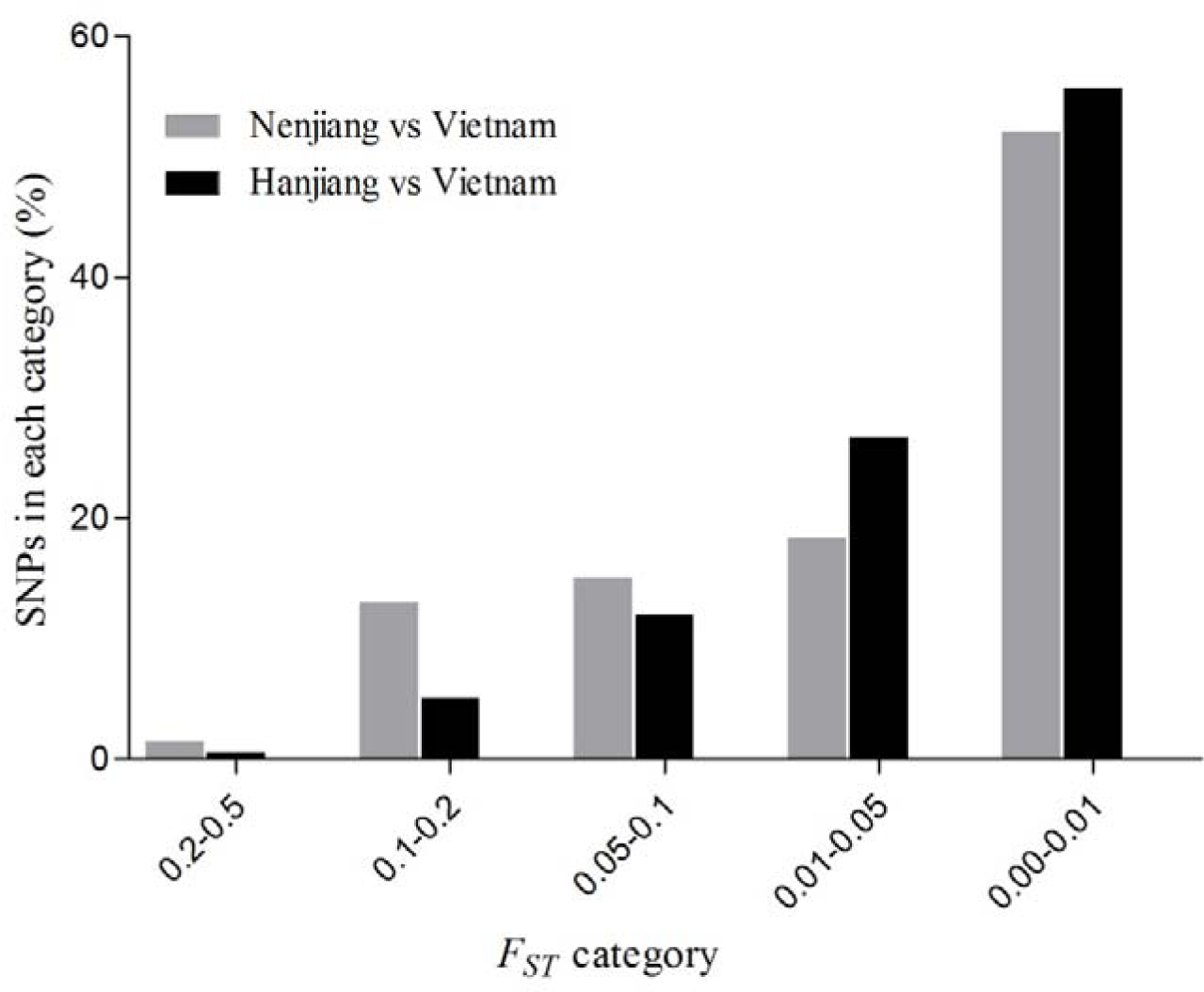
Distribution of *F_ST_* values in different categories between Nenjiang and Vietnam, with the longest geographical distance, and between Hanjiang and Vietnam, with the largest genetic distance, based on all genotyped SNPs with MAF > 0.05.

**Figure 3.**
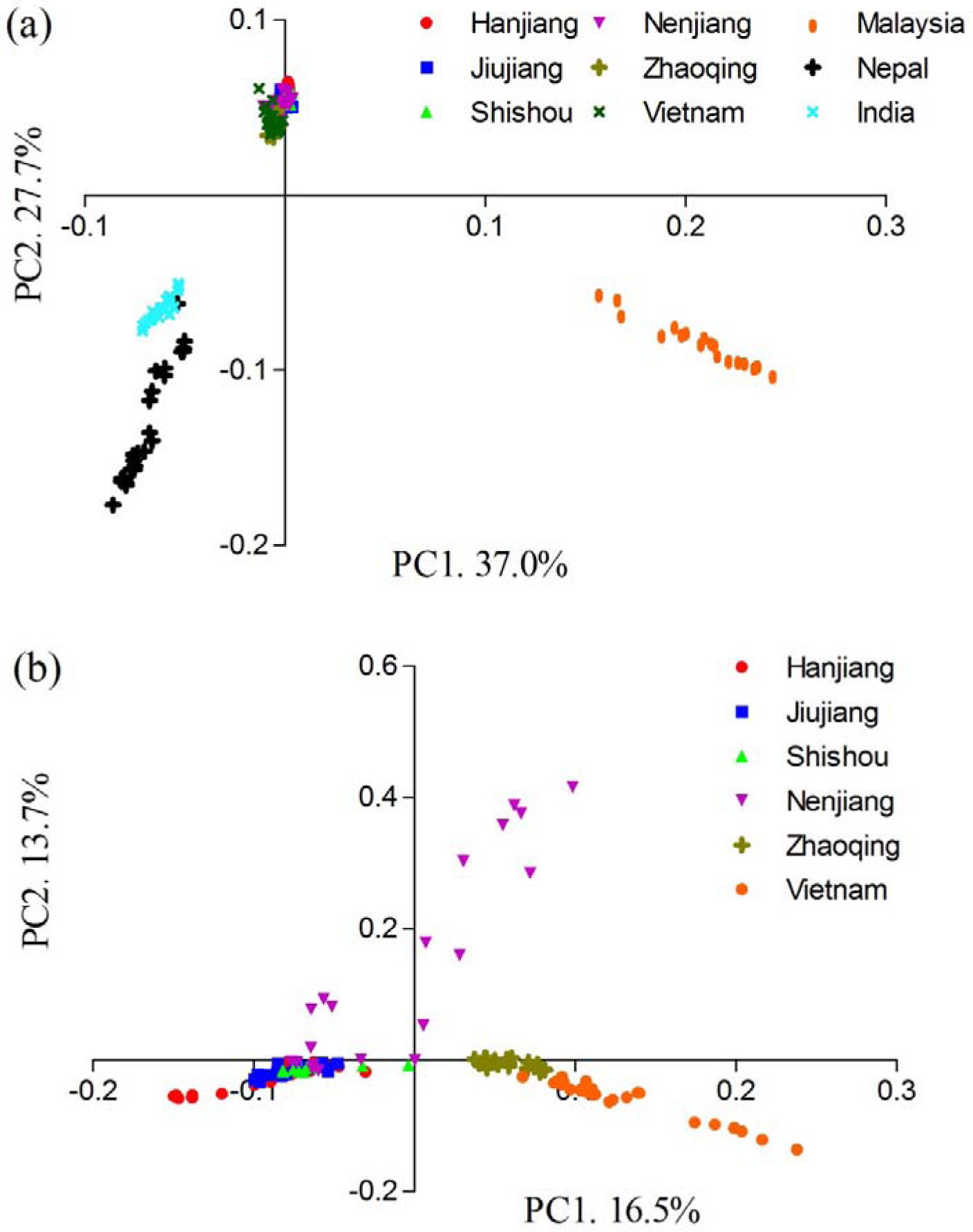
Principal component analyses for (a) all nine populations and (b) six native populations of grass carp based on all genotyped SNPs.

**Table 2.** Pairwise *F_ST_* values among each pair of populations of grass carp. Genetic differentiation that was non-significant after Bonferroni corrections (*P* < 0.001) is denoted in bold.

|  | Nenjiang | Hanjiang | Jiujiang | Shishou | Zhaoqing | Vietnam | India | Nepal | Malaysia |
| --- | --- | --- | --- | --- | --- | --- | --- | --- | --- |
| Nenjiang | - |  |  |  |  |  |  |  |  |
| Hanjiang | 0.0260 | - |  |  |  |  |  |  |  |
| Jiujiang | 0.0228 | 0.0154 | - |  |  |  |  |  |  |
| Shishou | 0.0196 | 0.0123 | **0.0073** | - |  |  |  |  |  |
| Zhaoqing | 0.0272 | 0.0351 | 0.0337 | 0.0279 | - |  |  |  |  |
| Vietnam | 0.0448 | 0.0515 | 0.0484 | 0.0449 | 0.0238 | - |  |  |  |
| India | 0.1265 | 0.1307 | 0.1282 | 0.1289 | 0.1126 | 0.1297 | - |  |  |
| Nepal | 0.1540 | 0.1580 | 0.1532 | 0.1596 | 0.1384 | 0.1592 | 0.1814 | - |  |
| Malaysia | 0.2295 | 0.2255 | 0.2224 | 0.2336 | 0.2205 | 0.2399 | 0.3167 | 0.3501 | - |

**Figure 4.**
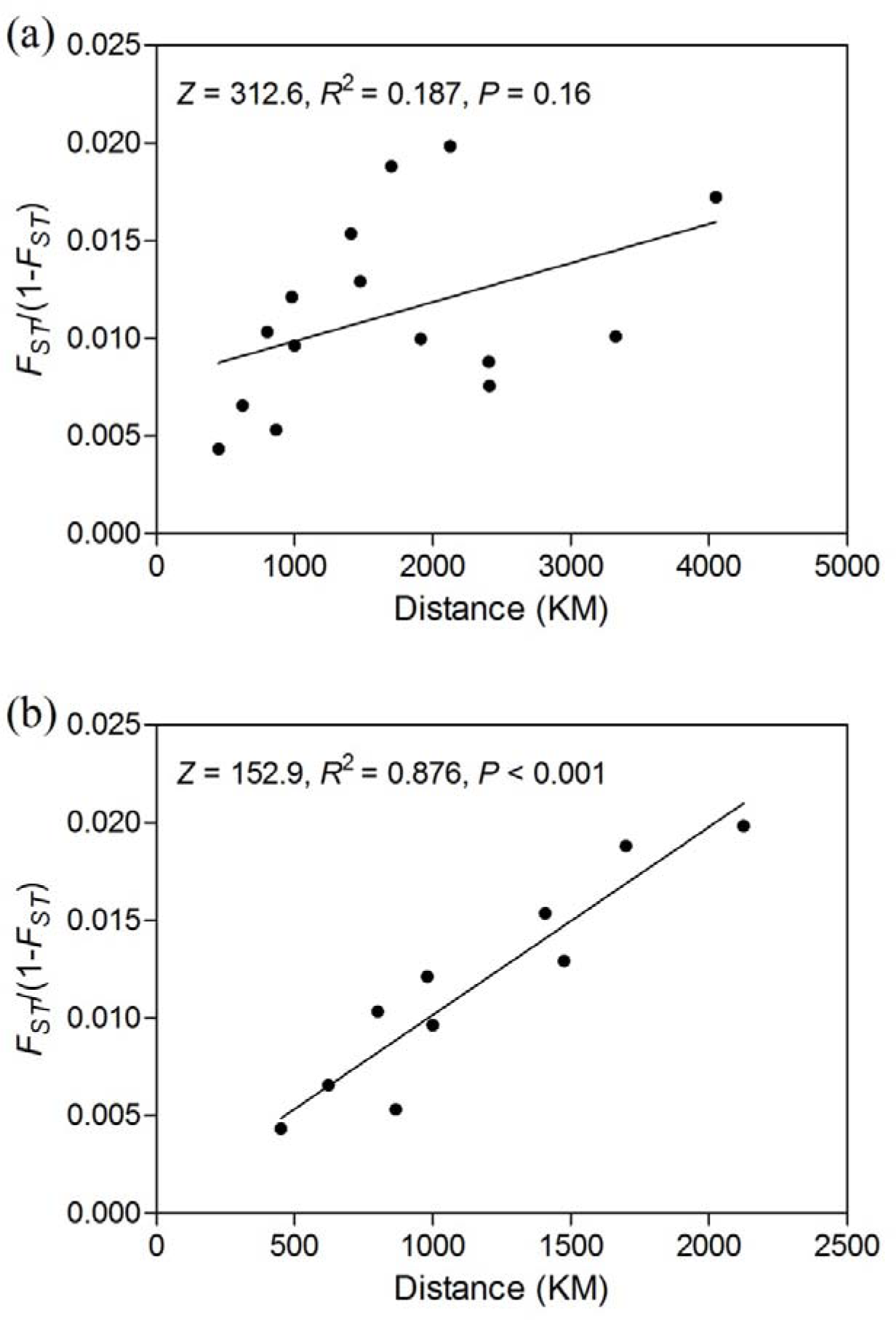
Figure 4 The overall pattern of isolation-by-distance for (a) all six native populations and (b) five native populations excluding population Nenjiang, examined using Mantel tests based on 5all genotyped SNPs. Genetic distance was estimated as *F_ST_*/(1-*F_ST_*), while geographical 6distance was the linear distance between sampling localities.

### Population introduction history

The phylogenetic tree showed that native populations from the Yangtze River (Hanjiang, Jiujiang and Shishou), the Heilongjiang River (Nenjiang) and the Pearl River (Zhaoqing and Vietnam) Systems formed three independent clusters, respectively, with the Heilongjiang River System cluster located between the Yangtze River System and the Pearl River System clusters (Figure 5a). For the introduced populations, India and Nepal formed one subcluster and joined into the Pearl River System cluster. Although joined into the Pearl River System, the introduced population Malaysia showed a relatively long genetic distance with the other populations within this cluster and was relatively close to the Heilongjiang River System. Considering the results of the principle component analysis, population Nenjiang and Malaysia might have an origin of admixture between the Yangtze River and the Pearl River System lineages (Figure 5a). We further inferred the origin of the two populations separately using ancestral alleles with the program fastStructure. Two populations of the closest genetic relationship, Jiujiang and Zhaoqing from the Yangtze River and the Pearl River System, respectively, were selected as the potential ancestral populations. Considering most of the SNPs have a very low level of genetic divergence (Figure 2), only loci of *F_ST_* > 0.05, numbering 5986, were used for these analyses. The most likely K values for estimation of the origins of the populations Nenjiang and Malaysia were inferred as 2 and 3, respectively (online supporting Figure S1). We found that many more ancestral alleles in the Nenjiang population in the Heilongjiang River System originated from the Yangtze River System than from the Pearl River System (Figure 5b). However, for the introduced population Malaysia, only ancestral alleles from the Pearl River System (Zhaoqing) were observed (Figure 5c).

**Figure S1.**
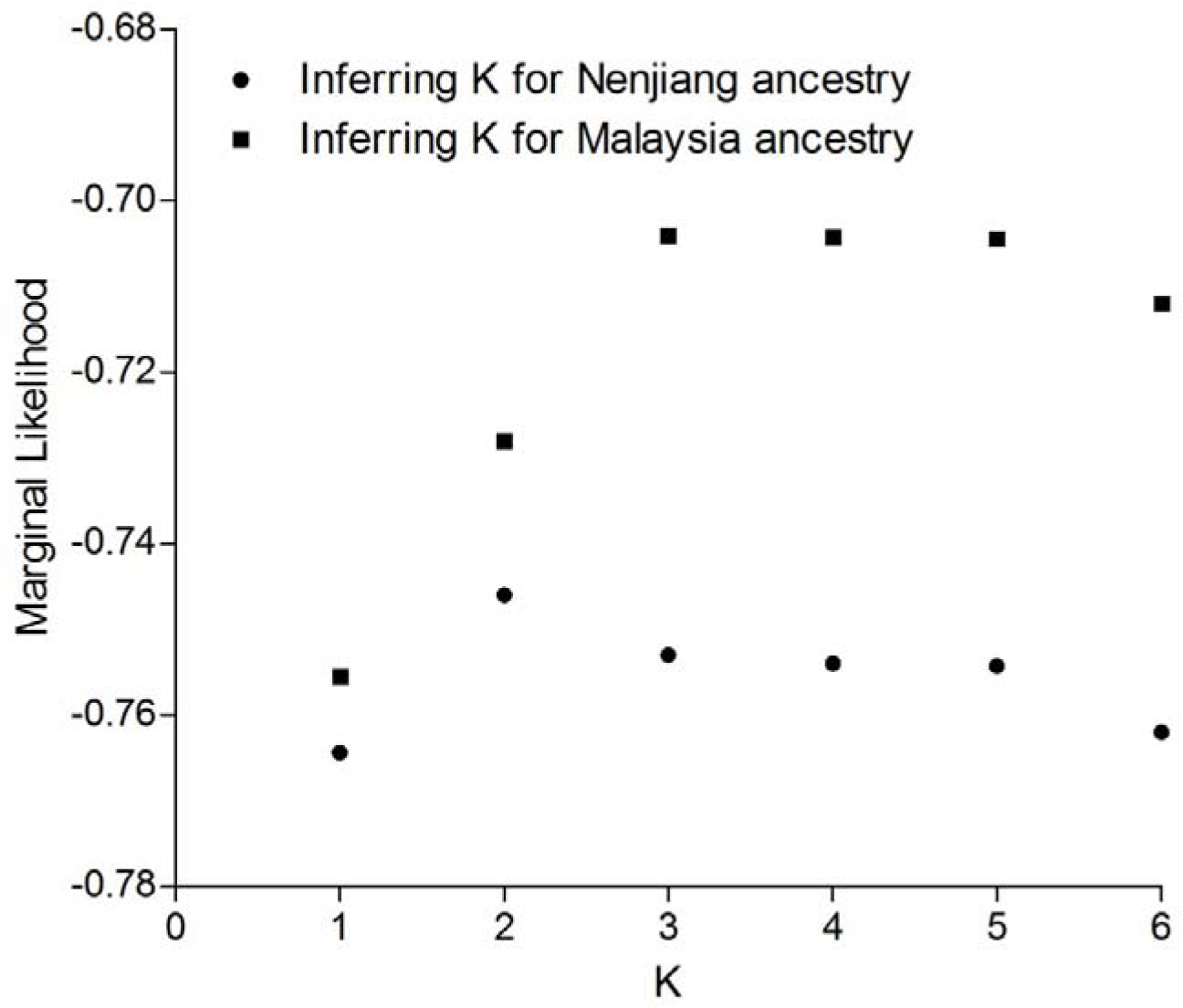
Plotting of K values inferred from the program fastStructure.

**Figure 5.**
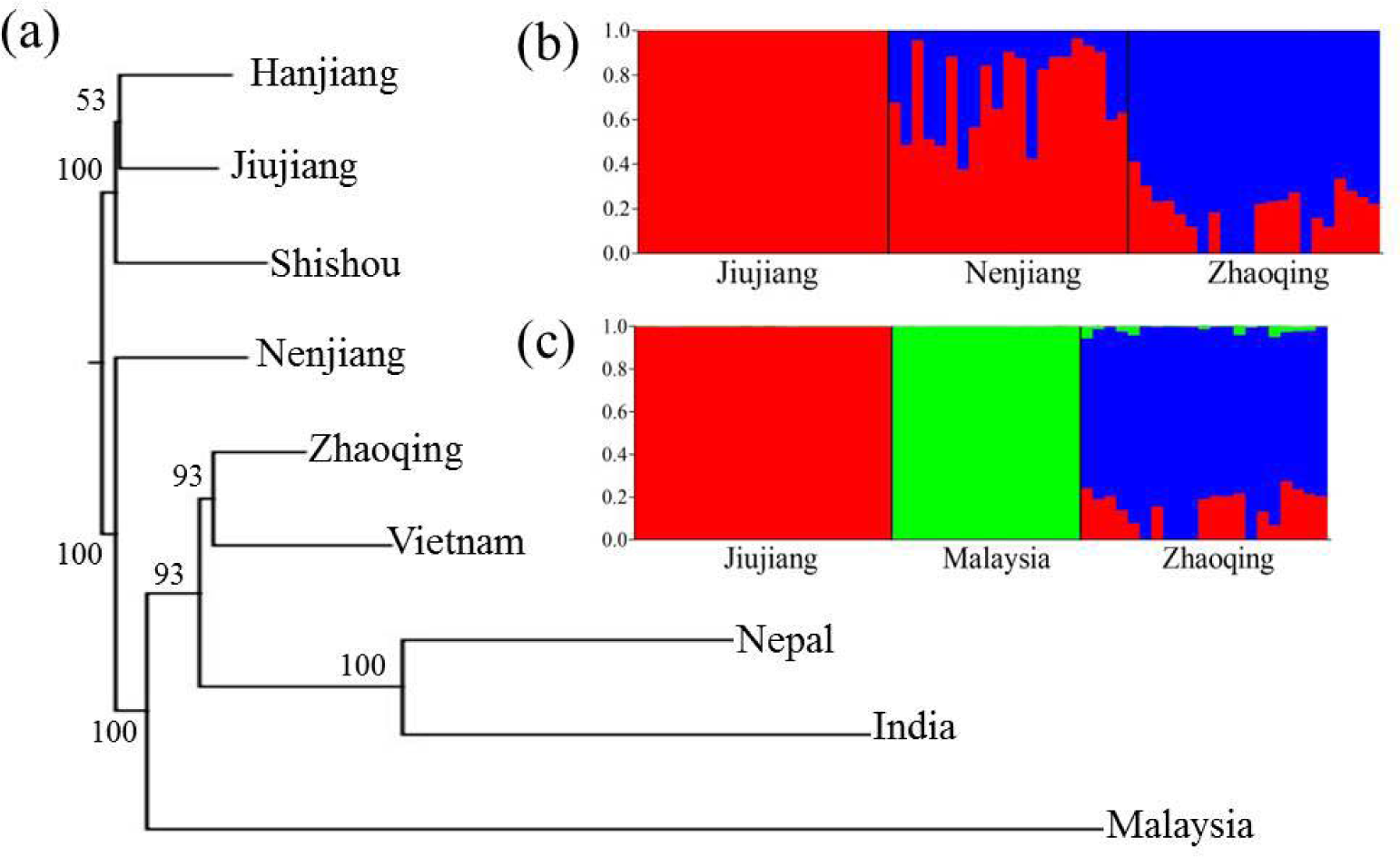
(a) Phylogenetic relationships among all nine populations of grass carp constructed using the Neighbor-Joining approach. Bootstrap supports over loci for 1000 times are indicated. (b) Genetic assignment of the native population Nenjiang to the Yangtze River System and the Pearl River System and (c) genetic assignment of the introduced population Malaysia to the Yangtze River System and the Pearl River System, respectively. The most likely K value for both assignment tests in the program Structure was inferred as 5. Each vertical line represents one individual, while each colour shows the genetic composition that is assigned into a distinct genetic cluster.

### Identifying loci under putative selection

Among all the SNPs with MAF > 0.05, 5197 (14.4%) were found to have significant linear regression (P<0.05) between latitudinal gradients and allele frequencies. Mantel tests revealed that 3351 (9.4%) SNPs showed significant patterns of isolation-by-distance (*R*^2^ > 0.514, *P* < 0.05) for individual locus. As stated above, the Nenjiang population showed a signature of admixture, and this produced decisive effects on the pattern of isolation-by-distance (Figure 3 & Figure 4). Therefore, we further removed this population from Mantel tests. Using five native populations, 3710 (10.4%) SNPs were observed to have significant patterns of isolation-by-distance (*R*^2^ > 0.632, *P* < 0.05). After removing loci with any evidence of isolation-by-distance, a total of 2700 loci that showed significant latitudinal variation in allele frequencies were obtained for further analysis (Figure 6).

**Figure 6.**
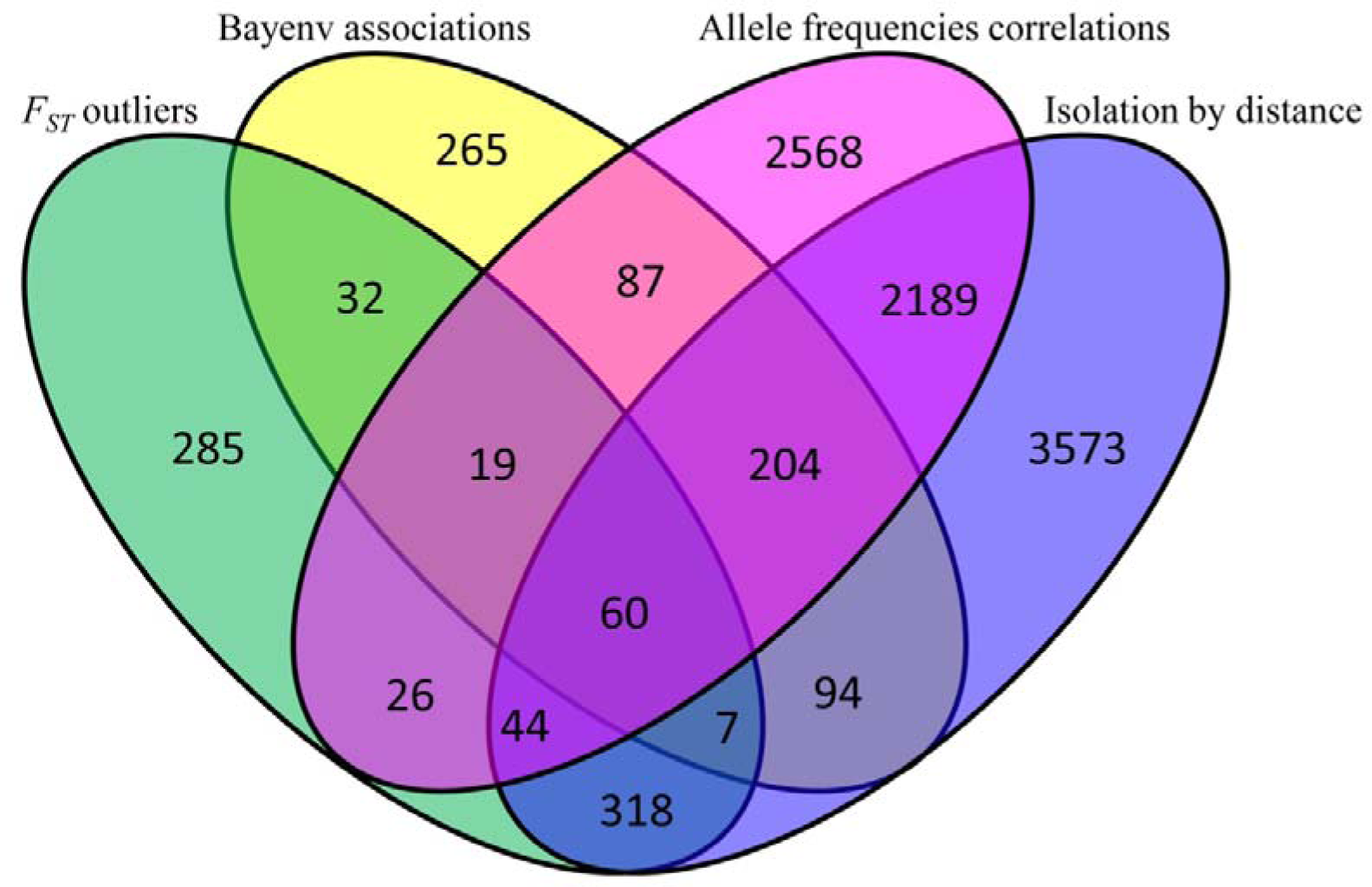
Venn diagrams showing the number of loci under putative selection and isolation-by-distance as revealed by *F_ST_* outlier tests, Bayenv association tests, Allele frequency correlations and Mantels tests for Isolation-by-distance. The numbers of overlapping loci among the different approaches are also illustrated.

The Bayesian generalized linear mixed model identified 768 SNPs that showed significant association between genetic variations and latitudinal gradients across all six populations at individual locus (BF > 3). In *F_ST_* based outlier tests, BayeScan detected 263 SNPs as substantial outliers (BF > 3), while the hierarchical island model identified 744 SNPs as significant outliers at the significance level of 0.99. A total of 791 (2.2%) unique loci were revealed to be outliers by the two *F_ST_* based tests.

Within the 2700 loci that showed correlation to latitudinal gradients, 132 were candidates under directional selection as revealed by the program Bayenv and both *F_ST_* based outlier tests (Figure 6). Moreover, Bayenv and the outlier tests identified 265 and 285 unique candidates under spatially purifying selection, respectively (Figure 6). We observed that the loci under putative selection revealed by the two outlier tests and Bayenv had much higher genetic divergence with mean *F_ST_* values of 0.198 and 0.104, respectively. However, the loci correlated to latitudinal gradients showed a much lower mean *F_ST_* (0.034) than the loci with the pattern of IBD (*F_ST_*, 0.065) and also the whole dataset (*F_ST_*, 0.037) (Figure 7). Low genetic divergence much likely suggests that selection pressure is weak on these loci. In order to reduce false positives, only loci that were revealed to be under potentially directional selection by outlier tests, Bayenv or allele frequencies association study, and also showed *F_ST_* of more than the 95% quantile (0.121) of the whole dataset, were retained, which produced 451 loci for further analyses.

**Figure 7.**
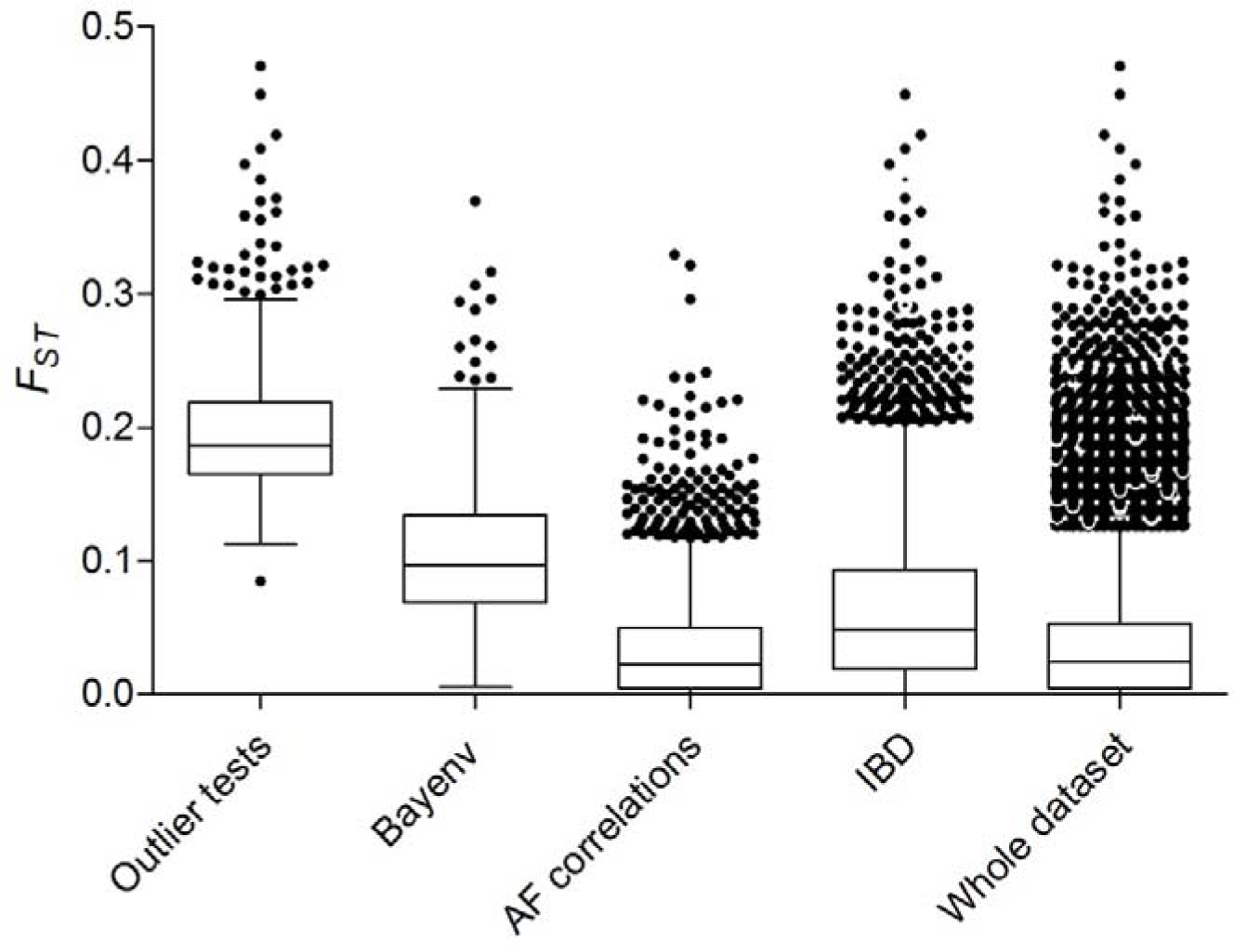
Distribution of *F_ST_* values for the SNPs that were identified as outliers for putative directional selection, associated with latitude as revealed by Bayenv, had allele frequencies correlated to latitude and agreed with the pattern of isolation-by-distance, as well as for the whole dataset. The highest and lowest error bar indicates the 95% quantile, while the median 1horizontal line denotes the mean *F_ST_*_value. Individual locus with F_ ST over the upper 95% 2quantile is shown.

### Functional annotation of genes under putative selection

Among the 451 loci, 84 (18.6%) were annotated as having a gene feature and were further investigated (Table S1). 42.9% (36) of the annotated genes were indicated to be significantly associated with latitudinal gradients as revealed by Bayenv or allele frequency correlation study. GO enrichment revealed that these genes covered a wide range of functions in biological processes: biological regulation, cellular process, developmental process, immune system process, metabolic process, pigmentation and response to stimulus (online supporting Figure S2). Three KEGG pathways: Focal adhesion, Vascular smooth muscle contraction and the Toll-like receptor signaling pathway, were enriched with each containing two genes (Table S2). Interestingly, by searching for literature, we found 16 (19.0%) genes that play important roles in immune responses (e.g. MHC I UAK and MHC II DAB) and/or antiviral responses (e.g. Myxovirus resistance proteins and Mitochondrial antiviral signalling protein) (Table 3). Among these immune-related genes, 8 (50.0%) showed a pattern of latitudinal variation as revealed by Bayenv or allele frequency correlation study, while the other 8 genes were suggested to be under spatially purifying selection (Table 3).

**Table 3.** Enriched candidate genes under putative directional selection and their potential functions in grass carp. Approaches, including *F_ST_* outlier tests, Bayenv and allele frequency correlations (AFC), that were used to determine if one gene was under putative selection, are also indicated.

| Locus | Gene name | Function | Approaches |  |  |
| --- | --- | --- | --- | --- | --- |
| Lca7796 | Toll-like receptor 5b | Immune response | $F_{ST}$ outlier | | |
| Lca7927 | Chemokine (C-C motif)-like | Immune response |  |  | AFC |
| Lca15228 | DEXH (Asp-Glu-X-His) box polypeptide 58 | Antiviral signaling | $F_{ST}$ outlier | | |
| Lca33879 | Phosphofurin acidic cluster sorting protein 2 | Antiviral signaling |  | Bayenv |  |
| Lca52319 | Myxovirus (influenza virus) resistance E | Antiviral signaling | $F_{ST}$ outlier | Bayenv | AFC |
| Lca57825 | Integrin alpha FG-GAP repeat containing 1 | Antiviral signaling |  | Bayenv | AFC |
| Lca79226 | Interleukin-1 receptor-associated kinase 1 | Immune response | $F_{ST}$ outlier | | |
| Lca88973 | Major histocompatibility complex class I antigen UKA | Immune response | $F_{ST}$ outlier | Bayenv | AFC |
| Lca110035 | Grass carp reovirus (GCRV)-induced gene 21 | Antiviral signaling | $F_{ST}$ outlier | | |
| Lca152085 | Interferon induced with helicase C domain 1 | Immune response | $F_{ST}$ outlier | | |
| Lca152284 | PC4 and SFRS1 interacting protein 1 | Antiviral signaling | $F_{ST}$ outlier | | AFC |
| Lca159757 | Major histocompatibility complex class II DAB | Immune response | $F_{ST}$ outlier | | |
| Lca175136 | Lymphocyte cytosolic protein 1 | Immune response |  |  | AFC |
| Lca183026 | Myxovirus (influenza) resistance A | Antiviral signaling | $F_{ST}$ outlier | | |
| Lca202544 | Baculoviral IAP repeat containing 2 | Immune response | $F_{ST}$ outlier | | |
| Lca285612 | Mitochondrial antiviral signalling protein | Antiviral signaling |  |  | AFC |

**Figure S2.**
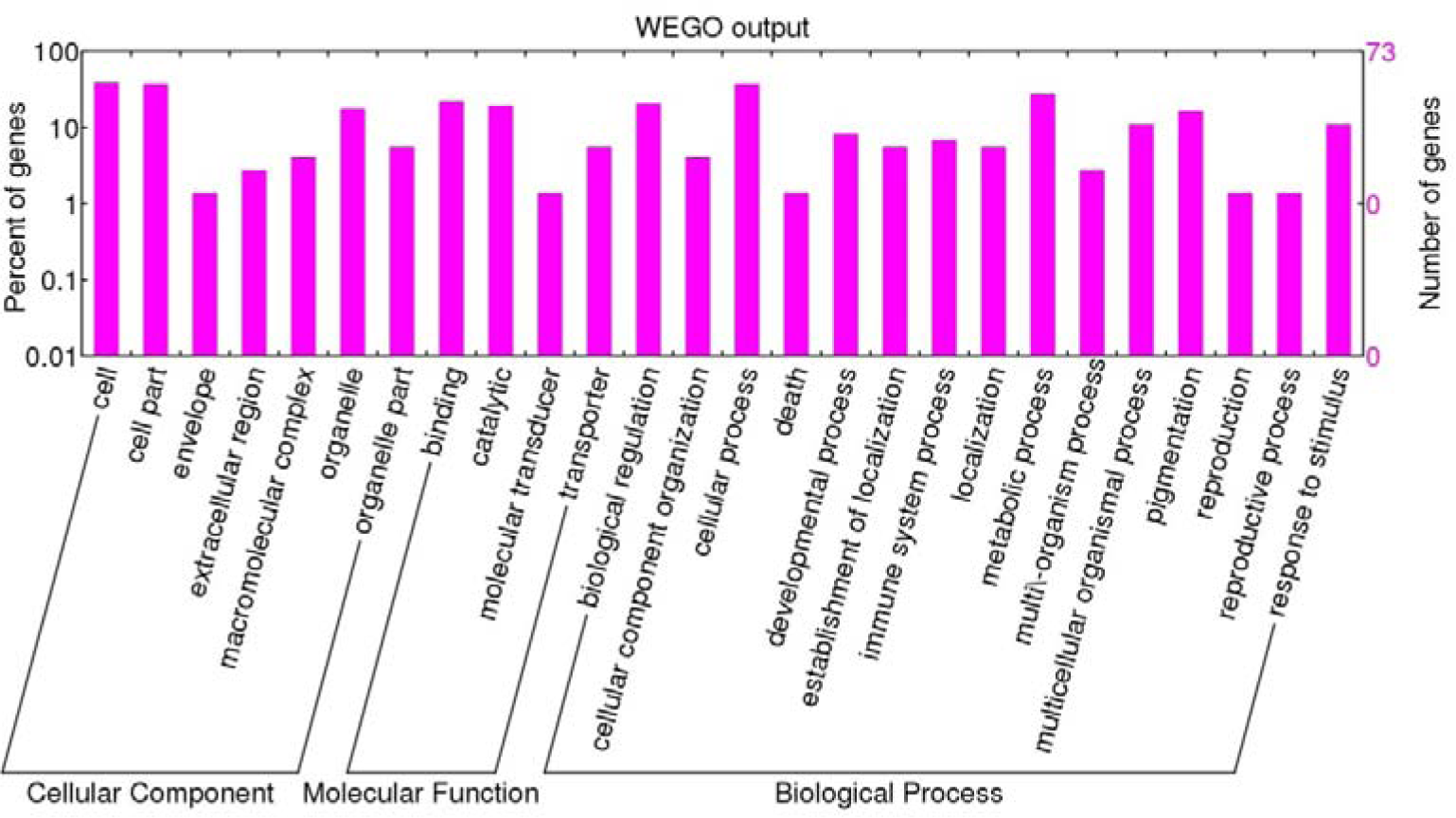
Gene ontology annotations of the candidate genes under potential local selection in grass carp. Three categories: Cellular Component, Molecular Function and Biological Process, were used to visualize the potential functions of enriched genes

## Discussion

In this study, we investigated range-wide population structure of native populations and origin of introduced populations in the South and Southeast Asia, as well as latitudinal variation and local selection of grass carp using population genomic approaches. This study provides important implications for future application of grass carp in the biocontrol of aquatic vegetation and understanding the mechanism of local adaptation, particularly adaptive latitudinal variation for freshwater fish species with a wide range of geographical distribution.

## High gene flow among native populations

Typically, freshwater fishes are more isolated by various geographical factors than marine fishes and thus have a lower level of gene flow (Ward *et al.* 1994). In this study, we observed that pairwise population genetic differentiation was very low (*F_ST_*, 0.0073-0.0515), comparable to a previous study using microsatellites (Liu *et al.* 2009). This level of gene flow across the native populations of grass carp is much higher than the average for freshwater species (Gyllensten 1985; Ward *et al.* 1994; Cooke *et al.* 2012) and even higher than some fishes living in the open marine environment, e.g. the Chinook salmon (*Oncorhynchus tshawytscha*) (Larson *et al.* 2014), Atlantic salmon (*Salmo salar*) (Bourret *et al.* 2013) and Asian seabass (*Lates calcarifer*) (Wang *et al.* 2015a). Further investigation on genetic differentiation at individual locus revealed that only < 15 % of total loci showed *F_ST_* > 0.1 and no loci had *F_ST_* > 0.5. Considering the geographical isolation among the three river systems, such results are rather unexpected. The genetic distance between the Pearl River System (Vietnam) and the Yangtze River System (Hanjiang) was closer than between the Pearl River System (Vietnam) and the Heilongjiang River System (Nenjiang), although the latter showed a rather longer geographical distance. Nevertheless, we observed that the number of loci with *F_ST_* > 0.1 was much more between Nenjiang and Vietnam than between Hanjiang and Vietnam. Considering the long aquaculture history of more than 1300 years, these contradicting results strongly suggest that the high gene flow among the three river systems is not only naturally occurring but also induced by human activities. Interestingly, the observed pattern of population differentiation did not conform to isolation-by-distance across the whole data set. However, after removal of the Nenjiang population that was suggested to have an admixture origin between the Yangtze River and the Pearl River Systems, the remaining populations showed a strong signal of isolation-by-distance. This indicates that although human-induced gene flow might have played important roles in shaping the overall population structure of grass carp, it only showed overwhelming importance in the Heilongjiang River System.

According to historical records, grass carp was abundant in both the Yangtze River System and the Pearl River System, and was widely captured from the wild as seeds for aquaculture locally (FAO 2014). There was no practical need to introduce grass carp between the two river systems. On the other hand, it is reasonable that gene flow can be high between the two river systems because they partially overlap in geography (Zhu 1993). For these reasons, the migration occurred much more naturally and thus the population differentiation showed a strong pattern of isolation-by-distance. However, we cannot exclude the possibility that human activities played important roles in dispersal of grass carp. Such gene flow might be merely induced so randomly with no directional purpose that it has much less effects on shaping genetic structure than the natural gene flow.

In contrast to the Yangtze and the Pearl River systems, the distribution and culture of grass carp in the Heilongjiang River System have never been abundant nor considered as a major aquaculture practice according to both historical records and current official fishery statistics (FAO 2014; Liu and Li 2015). Geographically, the Heilongjiang River System is completely isolated with the other two river systems. On the other hand, the Nenjiang population was mixed with alleles originating from both the Yangtze River and the Pearl River Systems as inferred from fastStructure. Hence, the low genetic differentiation between this river system and the other river systems strongly indicates that human-induced dispersal played more important roles than natural introgression. In fact, grass carp in the Heilongjiang River System grow slower than in the other river systems due to low water temperature (Cui *et al.* 1995). Therefore, seeds from the other river systems, particularly from the Yangtze River system, were commonly introduced for aquaculture purposes because of geographical adjacency. This is the most likely explanation for the low genetic differentiation of grass carp between the Heilongjiang River System and the other river systems.

## Origins of the early introduced populations

Both in terms of genetic diversity and differentiation, we observed significant genetic heterogeneity between all of the native populations and the introduced populations including Malaysia, India and Nepal, suggesting significant founder effects in the introduction of these populations (Barton and Charlesworth 1984). It is likely that the introduction of grass carp was not initiated under planned programs or that not all the introduced fish can adapt to the new habitats. Genetic differentiation among the native populations is the basis to trace the origins of the introduced populations (Cornuet *et al.* 1999; Paetkau *et al.* 2004). As expected, we identified significant genetic differentiation and also a clear geographical pattern of population differentiation among the three river systems in a background of high gene flow. Native populations from the Heilongjiang River, the Yangtze River and the Pear River Systems were separately clustered into independent genetic lineages, although there was evidence of population admixture for the Heilongjiang River System. These results provided critical clues to trace back population origins. Both pairwise *F_ST_* and phylogenetic analyses indicated that all three introduced populations, Malaysia, India and Nepal originated from the Pearl River System, which was also supported by the data inferred from ancestral alleles.

First, the Pearl River System is geographically more adjacent to Malaysia, India and Nepal than the Yangtze River and the Heilongjiang River Systems. Therefore, it is reasonable that the Pearl River System was preferred as the source for introduction to these countries. In contrast, the Yangtze River and the Heilongjiang River Systems are not only distant from Southeast and the South Asia, but also isolated by various continental barriers, e.g. the Himalaya Mountains. It is a great challenge to introduce fish from these two river systems to Southeast and the South Asia. Most importantly, it was recorded that grass carp was first introduced into Malaysia from Southern China with the large-scale migration of Chinese people in the 1800s, although it is not clear which river system the Malaysia population originated from (Welcomme 1988). The introduction history could be inferred from the routes of Chinese migration during that time. As revealed by history studies, most of the Chinese people in Southeast Asia were from Guangdong and Fujian provinces (Pan 1999), which geographically overlap with the Pearl River System. Thus, the Malaysia population very likely has an origin in the Pearl River System, consistent with the results of the genetic data.

In total, our data suggest that the native populations might have accumulated enough genetic divergence for population origin assignment of the recently introduced populations of grass carp, e.g. the populations introduced to Europe, North America and also some Southern Hemisphere countries (Mitchell 1986). These results are very valuable for studying the production and physiological adaptation, as well as the living environments and habitat preferences, of both native and introduced populations. Such information can be referenced to construct comprehensive introduction plans in the future.

## Local selection and latitudinal variation

It is a great challenge to discriminate local selection from neutral processes for organisms that have experienced complicated demographic history. Neutral processes can generate the same marks on genomic architecture as local selection does (Storz 2002; Vasemägi 2006; Savolainen *et al.* 2011; Wang *et al.* 2013; McKown *et al.* 2014; Hornoy *et al.* 2015). In some cases, adaptive traits show a specific distribution pattern along specific environmental factors. If the estimates of neutral forces are coincidentally varying along the same environmental factors, the difficulty of disentangling the roles of adaptive driving forces would be greatly enhanced (Merilä and Crnokrak 2001; McKay and Latta 2002; Storz 2002). Under this condition, a single association test between an individual locus and an environmental factor is obviously not enough to determine if one locus has experienced spatially divergent selection, particularly in the background of genome-wide patterns of isolation-by-distance (Vasemägi 2006). Grass carp is such a species, which has a significant signature of latitudinal distribution. Thus, the adaptive traits might vary in parallel with the pattern of neutral processes along specific geographical gradients, like latitude. These evolutionary processes limited the potential to identify the molecular mechanism underlying adaptive evolution (McKay and Latta 2002; Chen *et al.* 2012). Here, we used conceptually different approaches to differentiate the footprints of local selection from the currents of neutral evolutionary processes (Hansen *et al.* 2010; Wang *et al.* 2013).

Our main purpose was to identify individual loci of higher genetic divergence than can be explained by random genetic drift and gene flow (Storz 2002). As discussed above, grass carp from different river systems very likely have unique demographic history. Grass carp originated from the Yangtze River System and expanded into the Pearl River and the Heilongjiang River Systems during the Pleistocene and Pliocene, respectively (Li and Fang 1990). Nevertheless, the contemporary population structure was significantly shaped by high levels of gene flow due to both natural and artificial factors. As the Pearl River System and the Heilongjiang River System cover the southernmost and the northernmost distribution ranges, respectively, such contrasting environments have likely posed strong selective pressure on the distribution of grass carp (Gardner and Latta 2006). Gene flow within the Pearl River and the Yangtze River Systems might be seldom influenced by human activities, as population differentiation still shows a significant pattern of isolation-by-distance. However, gene flow between the Heilongjiang River System and the other two river systems were profoundly influenced by recent human activities, which overall changed the geographical pattern of population differentiation such that the pattern of isolation-by-distance was no longer observed. Although influenced by human activities, the extreme northernmost environmental condition of the Heilongjiang River System can pose strong selective pressure on the introduced grass carp. Such a process of natural selection provides important clues to discriminate footprints of natural selection from genome-wide patterns of isolation-by-distance.

Consistent with the overall neutral evolutionary process, a large number of loci, 6489 (18.1%) of the total loci, were indicated to show a pattern of isolation-by-distance in genetic divergence. Although 14.4% (5197) were revealed to have significant correlations between latitudinal gradients and allele frequencies, some of them would be false positives because these loci also showed significant correlations between pairwise geographical distance and genetic divergence. In total, these results suggest a strong background of isolation-by-distance in the overall population differentiation of grass carp. After removing the loci which were potential false positives by application of a series of conceptually different approaches, only 451 loci were suggested to be under putative directional selection, accounting for 1.3% of the total loci. This ratio is much less than in previous studies (2.3%-10%) using fish species that showed weak or non-significant genome-wide patterns of isolation-by-distance, like Atlantic salmon (Bourret *et al.* 2013), Chinook salmon (Larson *et al.* 2014) and yellowfin tuna (*Thunnus albacares*) (Grewe *et al.* 2015). Such results likely suggest that it is much less efficient to identify loci under putative directional selection with a genome-wide pattern of isolation-by-distance (Beaumont and Nichols 1996). Among the loci under putative directional selection, 18.6% were revealed to be associated with functional genes. Interestingly, 42.9% (36) of the annotated genes were indicated to be significantly correlated to latitudinal gradients, indicating clinally adaptive divergence at these loci (Storz 2002; Vasemägi 2006; Chen *et al.* 2012). Although these genes are involved in various functions, we observed a significant cluster of genes (16, 19.0%) playing important roles in immune responses (e.g. MHC I UAK and MHC II DAB) (Benacerraf 1981) and/or antiviral responses (e.g. Myxovirus resistance proteins and Mitochondrial antiviral signalling protein) (Seth *et al.* 2005; Gao *et al.* 2011). Among these immune-related genes, 8 (50.0%) showed a pattern of latitudinal adaptive variation. This result further suggests that the distribution of grass carp spanning approximately 30 latitudinal degrees was also the consequence of clinal adaptation along latitude. In the case of grass carp, the annual average temperature was observed to be highly correlated to the latitude of the sampling sites (*R*^2^ = 0.992, *P* < 0.001). However, we did not find any evidence that the enriched genes were associated to thermal adaptation. This result might suggest that the latitudinal adaptive distribution of grass carp was not directly selected by the temperature. Because the enriched genes were observed to have functions in defense against various pathogens and the diversity of pathogens were strongly related to the environmental temperature (Cashdan 2001; Mitchell *et al.* 2005;Dionne *et al.* 2007), our results might suggest that the latitudinal adaptation or clinal adaption of grass carp along latitude was the consequence of selection by pathogens and indirectly by temperature of the habitats.

In total, the joint application of different approaches identified a promising set of loci that were under putative directional selection. Many of them have a pattern of latitudinal variations. The latitudinal distribution of grass carp likely has an adaptive genetic basis, although the underlying causes remain to be elucidated. Nevertheless, spatially purifying selection has played important roles in shaping the contemporary population structure of grass carp. Our data shed light on the genetic basis of local adaptation of grass carp with a large distribution range.

## Acknowledgements

We thank Mr. Narayan Prasad Pandit and Mr. Dang Hai Nguyen for sample collection from Vietnam, Indian and Nepal respectively. This research is supported by the National Key Technology R&D Program of China (2012BAD26B02) and the China Agriculture Research System (CARS-46-04) foundations.

## Authors’ contributions

YS, LW, JL and GHY conceived the study and finalized the manuscript. YS and LW designed the experiments. YS, JF and XX carried out the lab experiments. LW and YS performed bioinformatics, analysed the molecular data and drafted the manuscript. All authors have read and approved the final manuscript.

## Supporting information

Table S1 Summary statistics of annotated genes under putative directional selection and their potential functions in grass carp

Table S2 Enriched KEGG pathways and the candidate genes under potential selection in grass carp

